# AtaA as a Self-Immobilizing Scaffold for Enzymatically Active Surfaces through Oxidative Tyrosine Coupling

**DOI:** 10.64898/2026.09.30.755837

**Authors:** Yuki Yokoe, Shogo Yoshimoto, Kosuke Minamihata, Rio Sugimoto, Ayasa Nagatani, Atsuki Nara, Noriho Kamiya, Katsutoshi Hori, Masahito Ishikawa

## Abstract

Protein nanofibers hold great promise as genetically programmable scaffolds for constructing functional material interfaces. AtaA, a giant trimeric autotransporter adhesin from *Acinetobacter* sp. Tol 5, forms an elongated architecture from the bacterial cell surface, enabling the cells to adhere directly to a wide variety of abiotic surfaces. However, the potential use of AtaA as a scaffold with additional biological functions remains largely unexplored. In this study, we investigated the use of AtaA as a self-immobilizing scaffold for constructing enzymatically active surfaces via horseradish peroxidase (HRP)-catalyzed tyrosine coupling. We prepared a recombinant AtaA nanofiber fragment (NhNs-AtaA) and its derivative containing two tyrosine-containing loop structures (YL-NhNs-AtaA). Both recombinant proteins retained their adhesive properties. Treatment with HRP and H₂O₂ generated high-molecular-weight products from both proteins, and dityrosine fluorescence emission indicated the formation of tyrosine-derived covalent cross-links. Y-Loop insertion further increased the consumption of monomeric AtaA and promoted the formation of high-molecular-weight products. The removal of arginine, which was included during purification, further increased the loss of monomeric AtaA in both constructs, indicating that intermolecular protein association influenced productive AtaA cross-linking. NhNs-AtaA and YL-NhNs-AtaA were combined with Y-tagged bacterial alkaline phosphatase (BAP-Y). HRP-mediated coupling generated high-molecular-weight products consistent with the cross-linked AtaA-BAP-Y conjugates and enabled the retention of BAP activity on AtaA-coated surfaces without prior chemical modification. Although Y-Loop insertion enhanced polymerization in solution, it reduced AtaA adhesion and consequently lowered surface-bound phosphatase activity, revealing a trade-off between cross-linking reactivity and native adhesive function. These findings establish AtaA as a self-immobilizing protein nanofiber scaffold for constructing enzymatically active surfaces, and suggest the potential extension of this strategy to other AtaA-based interfaces.

**TOC:** 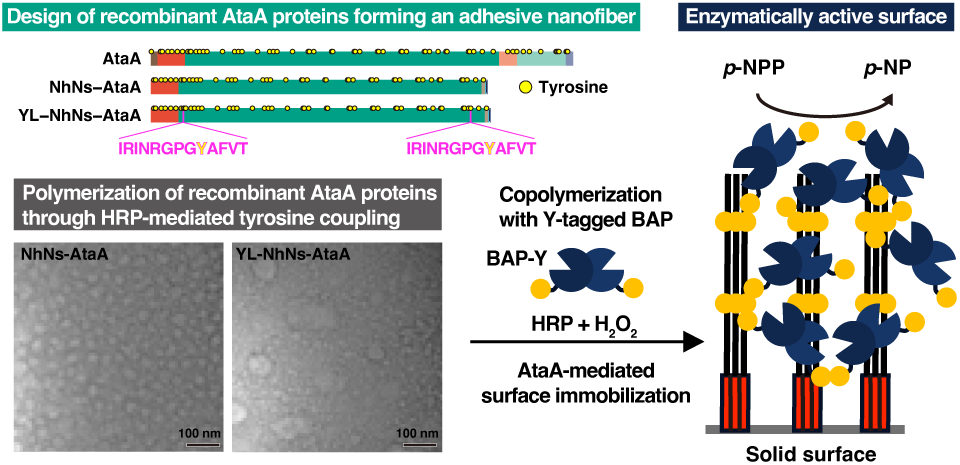

## Introduction

Proteins are genetically programmable building blocks of functional biomaterials. Unlike synthetic polymers, their molecular structures and functions are designed by editing the amino acid sequences encoded by DNA. This programmability enables the incorporation of diverse biological functions, including catalytic activity, molecular recognition, self-assembly, and adhesion, into well-defined molecular architectures ^1,2^. Among protein-based materials, fibrous proteins are particularly suited for constructing functional scaffolds because their elongated, repetitive architectures can organize molecular functions along extended protein assemblies. Recombinant spider silk proteins have been genetically functionalized with cell-adhesive motifs, such as fibronectin-derived or RGD-containing sequences, to generate silk-based materials that promote cell adhesion and growth on polymeric substrates ^3,4^. Engineered curli amyloid nanofibers have also been programmed to display functional peptide domains and are used as scaffolds for nanoparticle templating, substrate adhesion, covalent protein immobilization, and macroscopic nanofibrous materials ^5,6^. These examples highlight the potential of genetically encoded fibrous proteins as programmable scaffolds for the construction of functional materials.

AtaA, a giant trimeric autotransporter adhesin from the Gram-negative bacterium *Acinetobacter* sp. Tol 5, is a unique bacterial nanofiber protein whose adhesive function is encoded by its 3,630 amino acid sequence ^7–9^. AtaA forms an elongated fibrous architecture that extends from the bacterial cell surface, enabling cells to adhere strongly to a wide variety of abiotic surfaces ^10–13^. In addition to its adhesive properties, AtaA is recognized as a structurally robust nanofiber that retains its fibrous architecture under various conditions ^14–16^. As AtaA combines robust nonspecific adhesion with a structurally resilient nanoscale fibrous architecture, recombinant AtaA proteins may serve as useful scaffolds for constructing functional protein-based surfaces. However, strategies for functionalizing recombinant AtaA scaffolds with additional biological functions remain largely unexplored.

To convert programmable protein scaffolds into durable functional surfaces, the protein components must be stably organized into higher-order assemblies ^17–19^. Enzymatic tyrosine coupling offers a mild and genetically encodable strategy for this purpose, in which tyrosine-containing peptide motifs can be introduced into target proteins as defined reaction sites. In this reaction, accessible tyrosine residues are oxidized by oxidoreductases, such as horseradish peroxidase (HRP) and laccase, to form tyrosyl radicals, which subsequently couple to generate tyrosine-derived covalent cross-links, including dityrosine linkages ^20–22^. Previous studies introduced a tyrosine-containing peptide tag, the Y-tag, into target proteins and demonstrated site-selective cross-linking and polymerization of soluble proteins ^23–25^. In addition, a tyrosine-containing peptide loop, the Y-Loop, has been introduced into the loop region of bacterial alkaline phosphatase (BAP) to sterically control the coupling reaction, leading to the formation of linear protein polymers rather than irregular branched assemblies ^26^. Despite these advances, enzymatic tyrosine coupling has mainly been applied to soluble model proteins and enzymes, and its utility in the assembly of giant fibrous protein scaffolds remains largely unexplored.

In this study, we demonstrated the HRP-mediated covalent polymerization of recombinant AtaA nanofiber proteins through tyrosyl radical coupling. Recombinant AtaA was polymerized by HRP/H₂O₂, and dityrosine fluorescence supported the involvement of intrinsic tyrosine residues in the formation of covalent cross-links. The insertion of Y-Loops into recombinant AtaA further enhanced the polymerization efficiency, indicating that the engineered tyrosine-containing loops provided additional accessible coupling sites. Furthermore, this reaction can be extended to the copolymerization of AtaA with a Y-tagged enzyme, enabling enzyme accumulation on solid surfaces. This study expands enzymatic tyrosine-coupling polymerization from soluble model proteins to giant bacterial nanofiber proteins and provides a new strategy for constructing AtaA-based functional biomaterials.

## Results and Discussion

### Design and purification of AtaA recombinant proteins forming a nanofiber structure

Schematics of the wild-type AtaA and its recombinant proteins (NhNs-AtaA and YL-NhNs-AtaA) are shown in Figure 1A. Recombinant AtaA proteins were designed to be produced in the cytoplasm of *Escherichia coli* cells by removing the N-terminal signal peptide and transmembrane domain of the wild type. Their C-terminal regions were fused with the GCN4pII adaptor ^27^ to assist in the trimerization of the AtaA fragment ^15^ followed by His-tag and Strep-tag II. Although these affinity tags were expected to be used for protein purification, neither recombinant protein bound to their affinity resins. Similarly, we previously found that NhNs-AtaA-SNAP, the recombinant AtaA_59-2903_ fused to the GCN4pII adaptor, SNAP-tag, and Strep-Tag II at the C-terminus, does not bind to the affinity resin of Strep-tag II ^10^. The inefficient function of the C-terminal His- and Strep-tags is possibly due to their size. As Strep-Tag II (8 aa) and His-tag (6 aa) are much smaller than AtaA_59-2903_ (2,846 aa), the peptide tags may have difficulty approaching the affinity beads. Therefore, we adopted a loop-engineering strategy rather than terminal Y-tagging. Based on the strategy used in our previous study ^14^, in which a protease recognition sequence was inserted into a surface-exposed triglycine loop of an AtaA FGG domain, we introduced Y-Loops by extending analogous loop regions in FGG domains located near the N- and C-terminal regions of AtaA. This design was intended to place accessible tyrosine-coupling sites within the AtaA nanofiber scaffold and provide two engineered reaction sites per AtaA molecule. The primary structures of NhNs-AtaA and YL-NhNs-AtaA contained 48 and 50 tyrosine residues, respectively.

**Figure 1.**
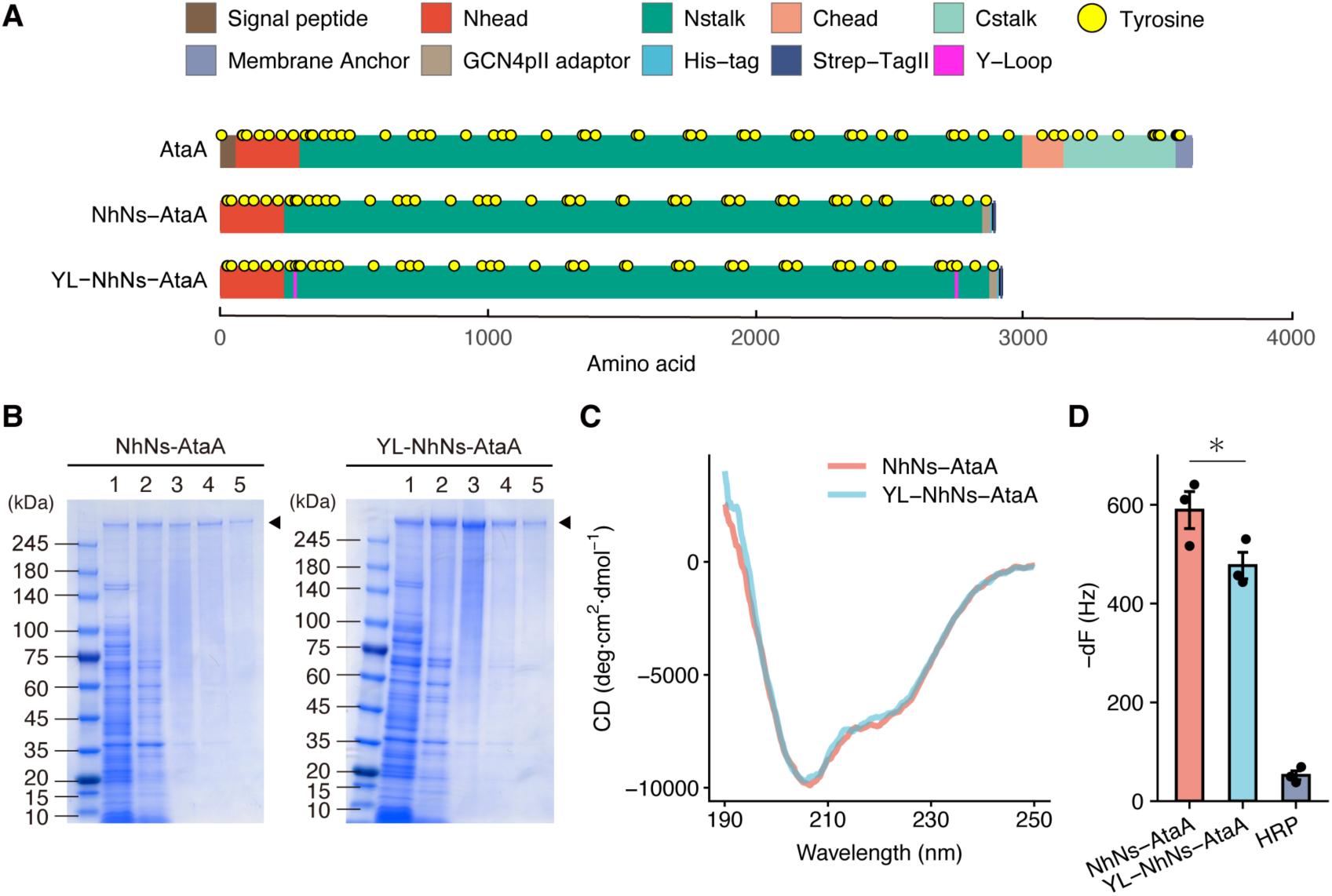
Preparation of recombinant AtaA proteins. (A) Schematics of native AtaA, NhNs-AtaA, and YL-NhNs-AtaA. (B) SDS-PAGE analyses of the purification process of NhNs-AtaA and YL-NhNs-AtaA. 1, cell lysate after sonication; 2, supernatant after the thermal treatment at 60 °C for 30 min; 3, pellet after 30% ammonium sulfate precipitation; 4, refolded protein fraction before the size exclusion chromatography; 5, purified protein fraction after size exclusion chromatography. Arrowheads indicate target proteins. (C) Circular dichroism spectra of purified NhNs-AtaA and YL-NhNs-AtaA proteins. (D) Quartz crystal microbalance analysis for the quantitative measurement of adhering recombinant proteins. Decrease in resonance frequency (−dF) after protein adsorption served as an index of the amount of protein retained on the surface. Horseradish peroxidase was included as a control. Bars represent the mean ± SEM of three technical replicates (n = 3). The asterisk indicates a significant difference according to an unpaired one-sided Welch’s *t*-test (*p* < 0.05).

Given the high molecular weight and thermostability of AtaA nanofibers ^14^, we developed a purification procedure based on heat treatment, ammonium sulfate precipitation, refolding, and size-exclusion chromatography (Figure 1B). The *E. coli* cell lysate, including the target proteins, was incubated at 60 °C for 30 min, and the precipitated proteins were removed by centrifugation. The remaining supernatant contained the target proteins and thermostable proteins derived from *E. coli* (lane 2 in Figure 1B). To remove low-molecular-weight proteins, 30% ammonium sulfate precipitation was performed. As the precipitate was not dissolved in any buffer not containing ammonium sulfate, it was dissolved in a 6 M guanidine solution (lane 3 in Figure 1B). Denatured proteins were refolded by dialysis against a buffer containing 200 mM arginine and subsequently separated by size exclusion chromatography (lanes 4 and 5 in Figure 1B). The secondary structures of the purified NhNs-AtaA and YL-NhNs-AtaA were analyzed by circular dichroism (CD) spectrometry. The CD spectra were comparable to those previously reported for folded recombinant AtaA proteins and were consistent with a mixed α-helical and β-strand-rich structure ^10,14^. To confirm whether the recombinant AtaA proteins inherited the adhesive function, we measured the adhesiveness of the purified proteins using a quartz crystal microbalance (QCM). Figure 1D shows the frequency change (dF) caused by the adhesion of the protein molecules to the QCM sensor when the frequency shift curve became mostly flat. The dF values of NhNs-AtaA and YL-NhNs-AtaA were much higher than that of HRP, a non-adhesive protein control, indicating the high adhesiveness of the AtaA recombinant proteins. Among the AtaA recombinant proteins, the dF value of YL-NhNs-AtaA was significantly lower than that of NhNs-AtaA (one-sided Welch’s *t*-test; *p* = 0.039). This difference suggests that the Y-Loop may slightly hamper the interaction of AtaA with surfaces.

### HRP-mediated polymerization of recombinant AtaA proteins via tyrosine coupling

To examine whether the purified recombinant AtaA proteins could be polymerized through oxidative tyrosine coupling, NhNs-AtaA and YL-NhNs-AtaA were incubated with HRP and H₂O₂. SDS-PAGE revealed that both proteins formed high-molecular-weight products near the top of the gel after treatment with HRP and H₂O₂, accompanied by a decrease in the bands corresponding to monomeric AtaA (Figure 2A). In contrast, no substantial change in electrophoretic mobility was observed when either HRP or H₂O₂ was omitted, indicating that the formation of high-molecular-weight products requires an HRP-catalyzed oxidative reaction. Despite lacking the introduced Y-Loops, NhNs-AtaA also formed a high-molecular-weight product. This suggests that endogenous tyrosine residues exposed on the surface of NhNs-AtaA can serve as substrates for HRP-mediated oxidation and intermolecular coupling. The 3–14% gradient gel used in this study can resolve proteins with molecular masses of up to approximately 1,500 kDa. The cross-linked products remained near the top of the gel, suggesting that their apparent molecular masses were beyond this separation range.

**Figure 2.**
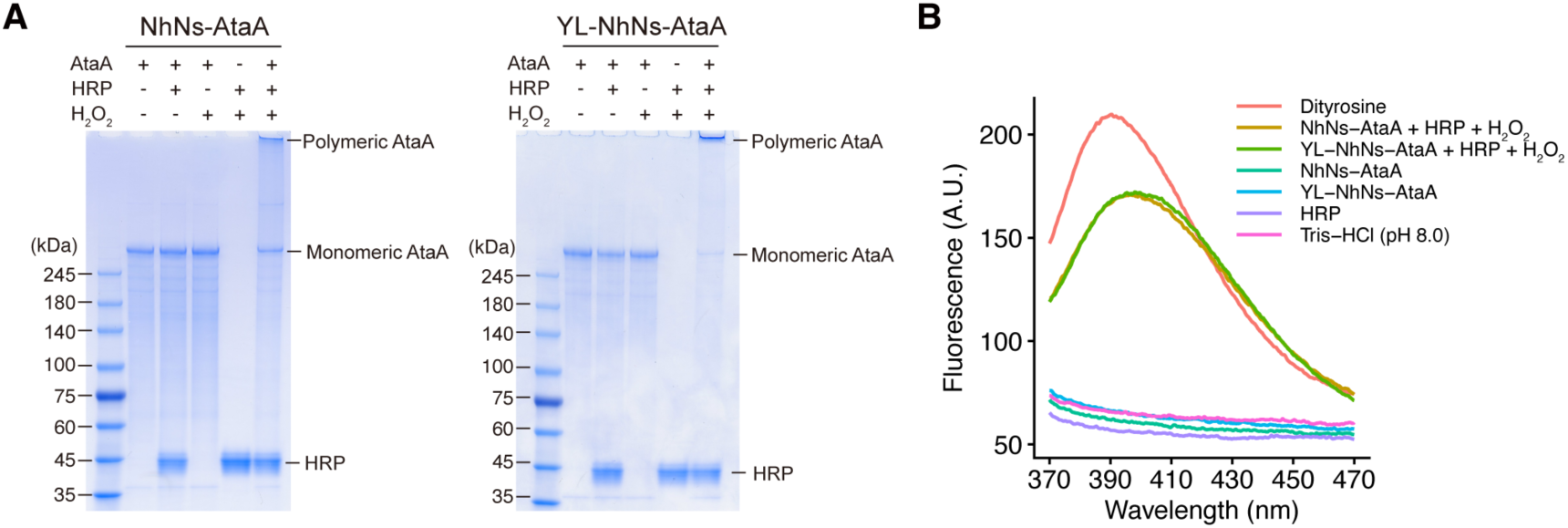
HRP-mediated polymerization of recombinant AtaA proteins. (A) SDS-PAGE analysis of NhNs-AtaA and YL-NhNs-AtaA incubated in the presence or absence of HRP and/or H₂O₂. (B) Fluorescence emission spectra of NhNs-AtaA and YL-NhNs-AtaA before and after incubation with horseradish peroxidase (HRP) and H₂O₂. The spectra were recorded upon excitation at 320 nm. HRP and dityrosine dissolved in 10 mM Tris-HCl buffer (pH 8.0) were measured as controls.

Given the apparent molecular mass of monomeric AtaA, this observation is consistent with assemblies of multiple AtaA molecules. The monomeric YL-NhNs-AtaA band appeared weaker than that of NhNs-AtaA after treatment with HRP and H₂O₂, suggesting that incorporation of the Y-Loops may have increased the efficiency of intermolecular cross-linking. Thus, although endogenous tyrosine residues were sufficient to support polymer formation, the introduced Y-Loop may have provided an additional accessible reaction site and promoted more efficient consumption of the monomeric protein.

To investigate whether polymerization involved tyrosine-derived cross-links, dityrosine fluorescence was examined. Dityrosine exhibits characteristic fluorescence at approximately 400–410 nm upon excitation at approximately 315–320 nm ^24^. HRP/H₂O₂-treated NhNs-AtaA and YL-NhNs-AtaA exhibited pronounced fluorescence with maxima at approximately 400 nm, whereas the untreated proteins, HRP alone, and buffer showed only weak fluorescence (Figure 2B). The emission profiles of the treated proteins were similar to that of the dityrosine standard, supporting the formation of dityrosine or related tyrosine-derived covalent cross-links during polymerization. Notably, the fluorescence spectra of HRP/H₂O₂-treated NhNs-AtaA and YL-NhNs-AtaA were nearly superimposable, suggesting that the overall formation of dityrosine-derived products may be comparable between the two proteins.

### Arginine suppresses intermolecular polymerization of recombinant AtaA proteins

Arginine hydrochloride was added at a concentration of 200 mM during purification of the recombinant AtaA proteins to maintain protein solubility. As arginine is widely used to suppress protein aggregation and protein–protein association ^28,29^, we hypothesized that residual arginine might also interfere with HRP-mediated intermolecular cross-linking of AtaA. To examine this possibility, arginine was removed from purified NhNs-AtaA and YL-NhNs-AtaA by dialysis, and the proteins were subsequently treated with HRP and H₂O₂. Three independently purified protein preparations were analyzed by SDS-PAGE (Figure 3A). Removal of arginine resulted in a greater loss of the monomeric AtaA band after HRP/H₂O₂ treatment for both recombinant proteins. This effect was particularly evident for YL-NhNs-AtaA, in which the monomeric band was barely detectable after arginine removal. To quantify these changes, the intensities of the monomeric AtaA bands were measured using ImageJ, and the percentage decrease after HRP/H₂O₂ treatment was calculated relative to the corresponding untreated sample (Figure 3B). Two-way mixed-design analysis of variance (ANOVA) revealed significant main effects of the protein construct (*F*(1, 4) = 322.28, *p* < 0.001) and arginine treatment (*F*(1, 4) = 155.58, *p* < 0.001), with no significant interaction between the two factors (*F*(1, 4) = 1.29, *p* = 0.320). This indicated that the Y-Loop introduction and arginine removal independently promoted the incorporation of AtaA molecules into high-molecular-weight products. Thus, the Y-Loop provides an effective reaction site for intermolecular tyrosine coupling, whereas arginine suppresses productive AtaA polymerization.

**Figure 3.**
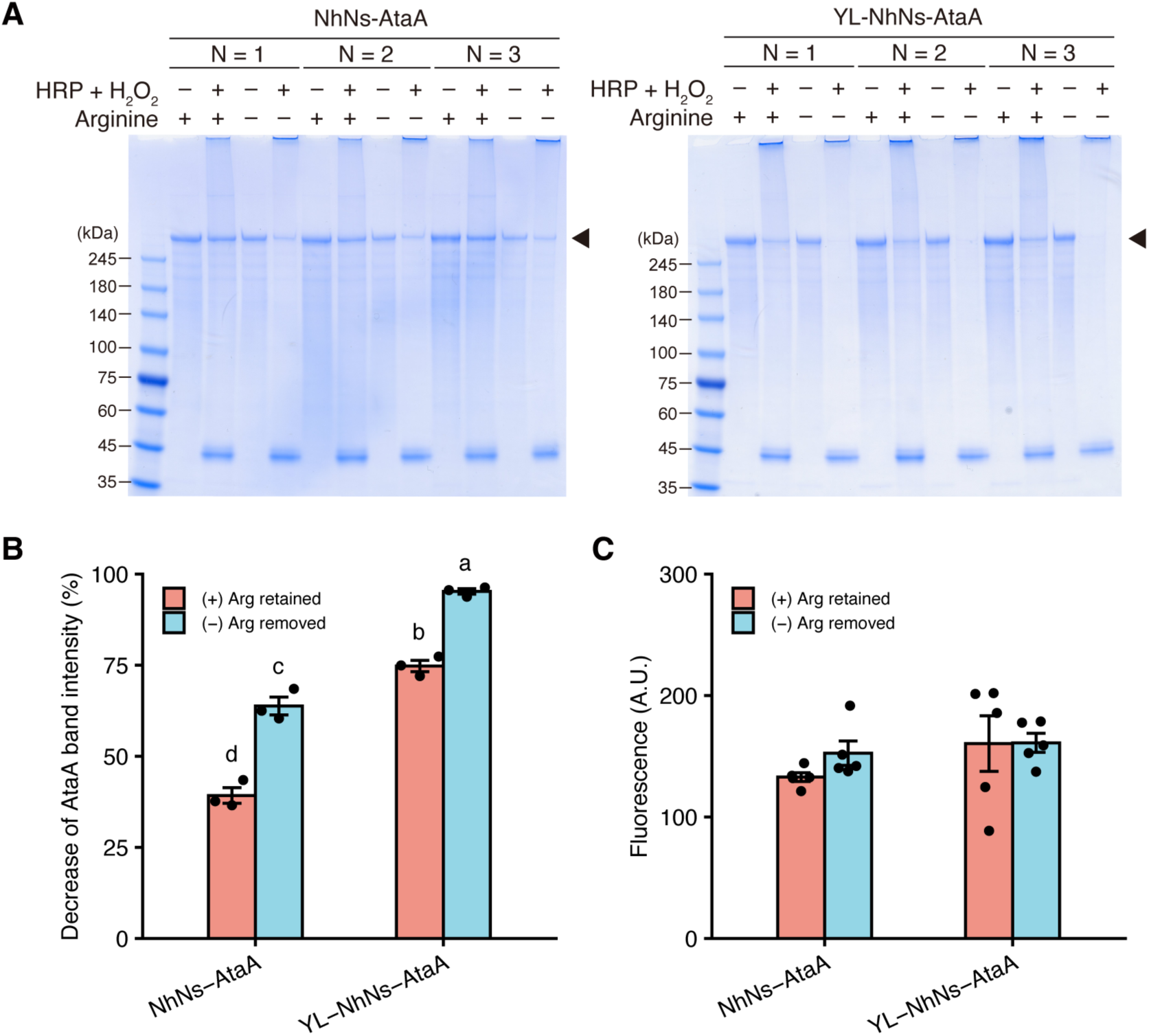
Effect of arginine on the horseradish peroxidase (HRP)-mediated cross-linking of AtaA recombinant proteins. (A) SDS-PAGE analysis of three independently purified preparations of NhNs-AtaA and YL-NhNs-AtaA after HRP/H₂O₂ treatment in the presence or absence of arginine. Arrowheads indicate monomeric AtaA. Arginine + indicates purified protein containing 200 mM arginine, whereas Arginine − indicates the corresponding protein preparation after dialysis against arginine-free buffer. (B) Densitometric quantification of the decrease in monomeric AtaA band intensity after HRP/H₂O₂ treatment. Arg retained and Arg removed correspond to the Arginine + and Arginine − conditions shown in panel A, respectively. Band intensities in the SDS-PAGE images shown in (A) were quantified using ImageJ, and the percentage decrease was calculated relative to the corresponding sample without HRP/H₂O₂ treatment. Bars represent the mean ± SEM of three independently purified protein preparations (n = 3). Individual data points are shown. Different lowercase letters indicate significant differences among the four conditions according to Tukey’s multiple-comparison test following two-way mixed-design analysis of variance (ANOVA) (*p* < 0.05). (C) Dityrosine fluorescence of NhNs-AtaA and YL-NhNs-AtaA after HRP/H₂O₂ treatment in the presence or absence of arginine. Bars represent the mean ± SEM of 5 independent reaction experiments performed on separate days (n = 5). Individual data points are shown.

Next, we measured dityrosine fluorescence in the reaction products to determine whether the enhanced disappearance of monomeric AtaA was accompanied by an increase in the overall formation of tyrosine-derived cross-links (Figure 3C). No statistically significant differences in fluorescence intensity were detected between the two protein constructs or between the presence and absence of arginine. The relatively large variation among independent reaction experiments may have limited the detection of modest differences. Nevertheless, the absence of a detectable increase in dityrosine fluorescence contrasts with the clear changes observed using SDS-PAGE. This discrepancy suggests that arginine removal primarily altered the balance between intramolecular and intermolecular tyrosine-derived cross-links rather than markedly increasing the total amount of fluorescent dityrosine products. Dityrosine can be formed through both intermolecular and intramolecular coupling between endogenous tyrosine residues. Therefore, in the presence of arginine, tyrosine coupling may have occurred without efficiently connecting separate AtaA molecules, whereas the removal of arginine may have increased the probability that separate AtaA molecules approached one another in a configuration favorable for intermolecular coupling, thereby promoting polymer formation. Similarly, the introduction of the Y-Loop may have provided an accessible cross-linking site that more efficiently converted tyrosine coupling events into intermolecular connections.

Arginine is widely used to suppress protein aggregation and association, although the underlying molecular mechanisms remain incompletely understood. Proposed mechanisms include direct interactions of the guanidinium group with aromatic and charged residues, as well as changes in protein solvation and protein–protein interactions ^28–30^. However, the fluorescence results do not support a major effect of arginine removal on the overall formation of tyrosyl radicals or dityrosine. Rather, arginine likely reduced the probability of short-lived tyrosyl radicals on separate AtaA molecules approaching each other sufficiently close to undergo intermolecular coupling. Together, these results suggest that productive molecular association is an important determinant of HRP-mediated AtaA polymerization and that removal of arginine allows the introduction of a Y-Loop to promote efficient intermolecular cross-linking.

### Transmission electron microscopy (TEM) of recombinant AtaA after HRP/H₂O₂ treatment under arginine-free conditions

To visualize the morphology of the reaction products, NhNs-AtaA and YL-NhNs-AtaA were examined by TEM after HRP/H₂O₂ treatment under arginine-free conditions. Both samples contained elongated fibrillar structures resembling the characteristic morphology previously reported for native AtaA fibers ^8,14^ (Figure 4). In addition to these fibrillar structures, non-fibrillar assemblies were observed in both samples. In the NhNs-AtaA sample, small non-fibrillar assemblies were observed among the fibrillar structures (Figures 4A and 4B), whereas the YL-NhNs-AtaA sample contained both small and markedly larger assemblies (Figures 4C and 4D). Representative fibrillar structures and large non-fibrillar assemblies are indicated by arrows and asterisks, respectively (Figures 4B and 4D). Such non-fibrillar assemblies were not evident in representative images of the corresponding proteins before HRP/H₂O₂ treatment (Figure S1). These observations suggest that HRP-mediated cross-linking promotes the formation of higher-order AtaA assemblies. Notably, the non-fibrillar assemblies observed for YL-NhNs-AtaA appeared larger and more extensive in the representative images than those observed for NhNs-AtaA. This morphological difference suggested that the introduced Y-Loop facilitated the incorporation of a greater number of AtaA molecules into intermolecularly cross-linked assemblies. This interpretation is consistent with the greater decrease in monomeric YL-NhNs-AtaA band intensity after HRP/H₂O₂ treatment. Although TEM does not directly distinguish covalent polymers from non-covalent aggregates, the observed morphological differences were consistent with the SDS-PAGE results.

**Figure 4.**
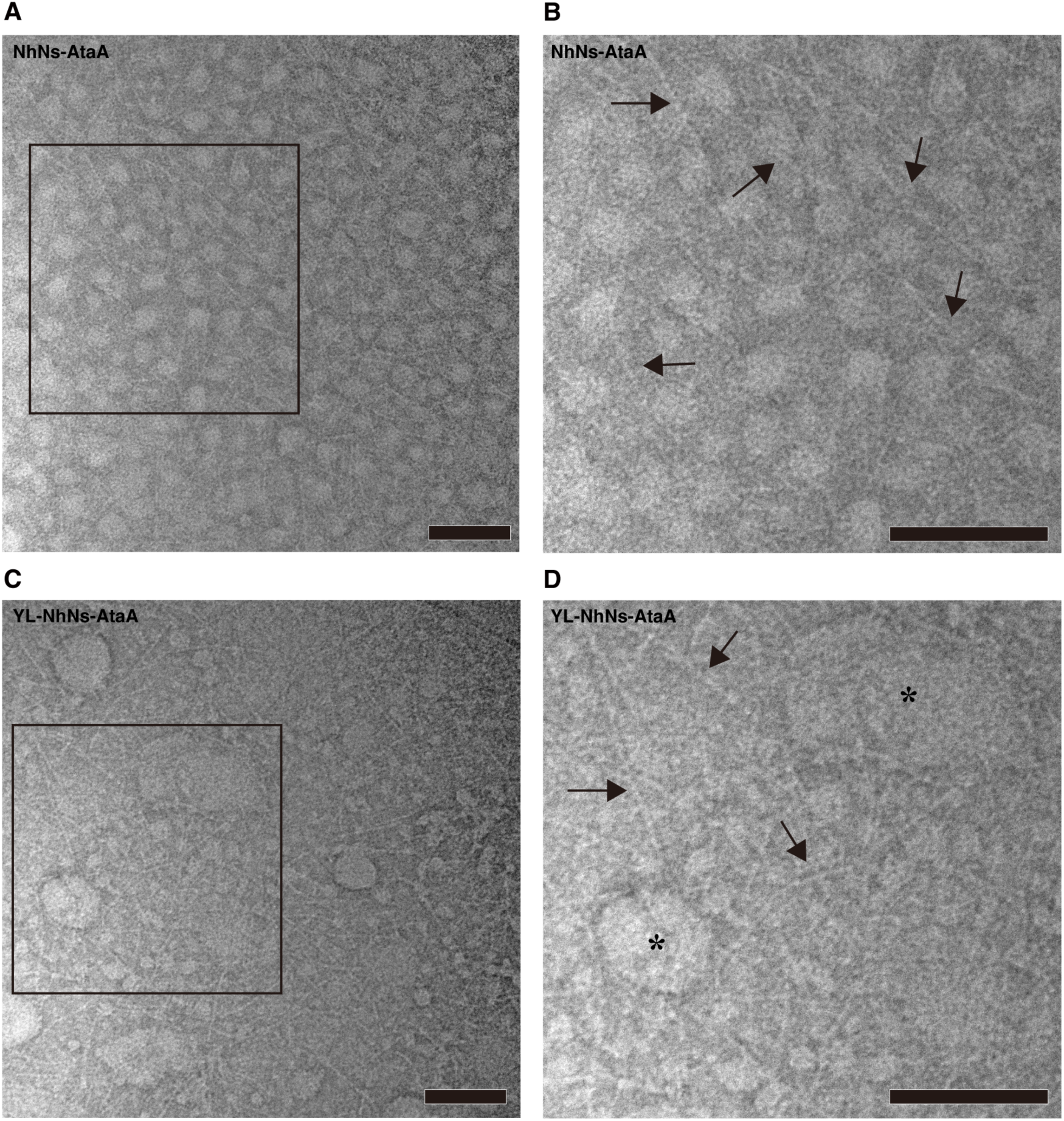
Transmission electron microscopy observation of horseradish peroxidase (HRP)-mediated AtaA polymers. (A, C) Representative transmission electron microscopy images of NhNs-AtaA (A) and YL-NhNs-AtaA (C) after HRP/H₂O₂ treatment in the absence of arginine. Boxed regions are shown at higher magnification in panels B and D, respectively. (B, D) Magnified views of the boxed regions in panels A and C. Arrows indicate representative elongated fibrillar structures, and asterisks indicate representative large non-fibrillar assemblies. Scale bars, 100 nm.

### AtaA-mediated construction of an enzymatically active surface

To examine whether the adhesive AtaA scaffold could be endowed with enzymatic activity through tyrosine coupling, NhNs-AtaA or YL-NhNs-AtaA was reacted with Y-tagged bacterial alkaline phosphatase (BAP-Y) in the presence of HRP and H₂O₂. SDS-PAGE analysis showed the formation of high-molecular-weight products near the top of the gel when AtaA and BAP-Y were reacted together in the presence of HRP and H₂O₂ (lanes 2, 4–6, 8, and 9 in Figure 5A). These products were not observed when HRP/H₂O₂ was omitted (lanes 1 and 3 in Figure 5A) or when AtaA was absent (lanes 7 and 10 in Figure 5A). The appearance of high-molecular-weight products, together with a decrease in the monomeric BAP-Y bands, suggested the formation of cross-linked conjugates containing both AtaA and BAP-Y. Increasing the BAP-Y:AtaA molar ratio from 1:1 to 2:1 resulted in a greater decrease in the monomeric and dimeric BAP-Y bands, suggesting that a larger amount of BAP-Y was incorporated into the AtaA-containing polymers (lanes 8 and 9 in Figure 5A). In contrast, BAP-Y formed a dimeric product under the same reaction conditions (lanes 7 and 10 in Figure 5A).

**Figure 5.**
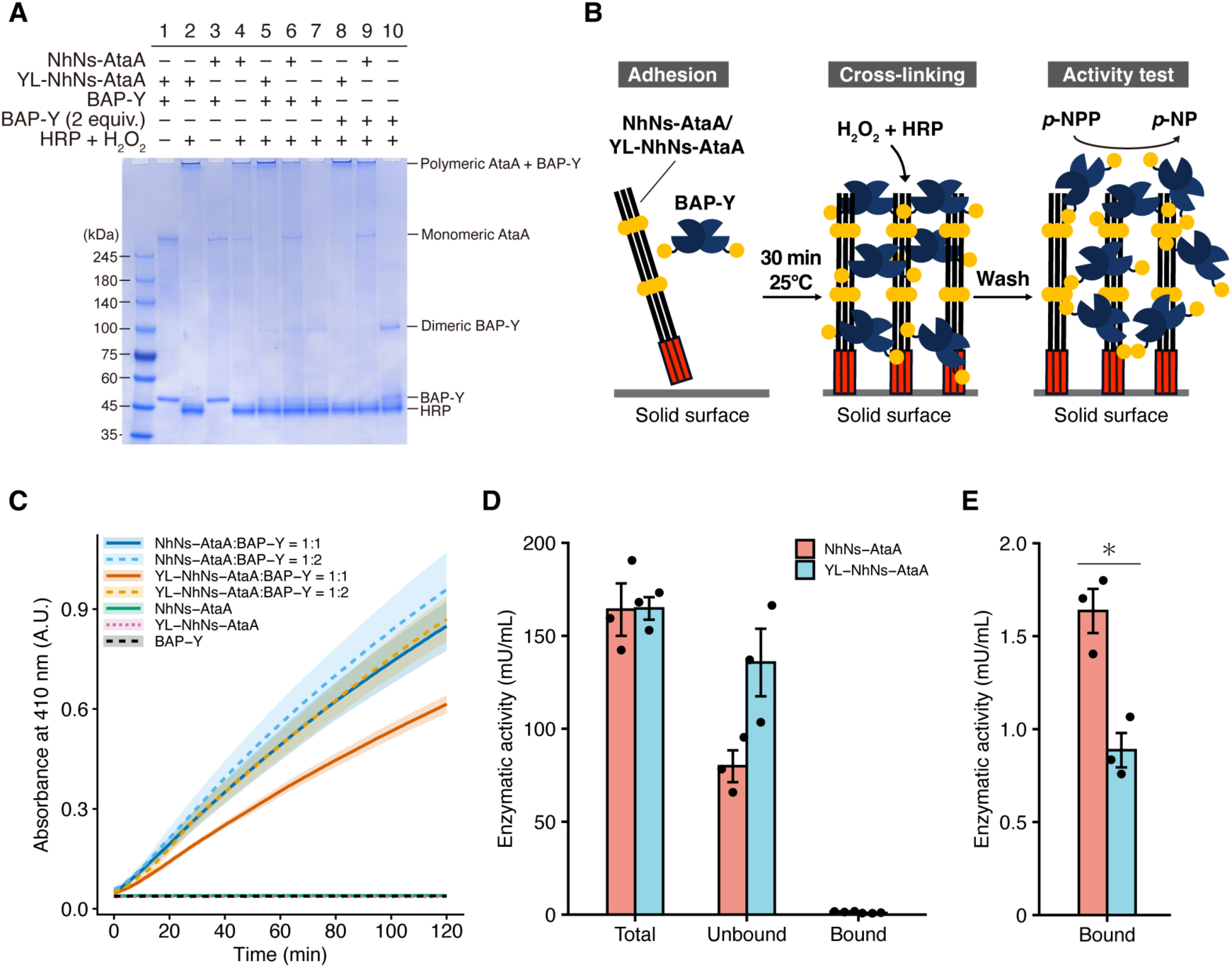
Construction of an enzymatically active surface through AtaA-mediated immobilization and horseradish peroxidase (HRP)-catalyzed cross-linking. (A) SDS-PAGE analysis of the HRP-mediated cross-linking of NhNs-AtaA or YL-NhNs-AtaA with Y-tagged bacterial alkaline phosphatase (BAP-Y). AtaA and BAP-Y were mixed at molar ratios of 1:1 or 1:2 and incubated in the presence or absence of HRP and H₂O₂ as indicated above the gel. BAP-Y without AtaA was analyzed under the same conditions. Positions of polymeric AtaA–BAP-Y products, monomeric AtaA, dimeric BAP-Y, monomeric BAP-Y, and HRP are indicated. (B) Schematic illustration of the construction and activity measurement of the enzymatically active surface. (C) Time courses of *p*-nitrophenol formation on surfaces treated with NhNs-AtaA or YL-NhNs-AtaA and BAP-Y at AtaA:BAP-Y molar ratios of 1:1 or 1:2. Surfaces treated with NhNs-AtaA alone, YL-NhNs-AtaA alone, or BAP-Y alone were used as controls. (D) Alkaline phosphatase activities in the total reaction mixture, unbound fraction, and surface-bound fraction after immobilization at an AtaA:BAP-Y molar ratio of 1:1. (E) Enlarged comparison of the surface-bound activities shown in (D). Data in (C–E) are presented as the mean ± SEM of three independent experiments. The asterisk indicates a significant difference according to an unpaired two-sided Welch’s *t*-test (*p* < 0.01).

Next, we constructed an enzymatically active surface using this cross-linking reaction, as shown in Figure 5B. NhNs-AtaA or YL-NhNs-AtaA was mixed with BAP-Y, and the mixture was incubated in the wells for 30 min at 25 °C to allow AtaA to adsorb onto the solid surface. HRP and H₂O₂ were then added to initiate covalent cross-linking between surface-associated AtaA and BAP-Y. After incubation for 1 h at 37 °C, the reaction mixture was removed, and the wells were washed to remove unbound proteins. The alkaline phosphatase activity retained on the surface was evaluated by directly adding *p*-nitrophenyl phosphate (*p*-NPP) and monitoring the formation of *p*-nitrophenol (*p*-NP) at 410 nm. A time-dependent increase in absorbance was observed when AtaA and BAP-Y were combined, whereas surfaces treated with NhNs-AtaA, YL-NhNs-AtaA, or BAP-Y alone showed little change in absorbance (Figure 5C). These results indicate that BAP-Y alone was not efficiently retained on the surface and that stable immobilization required both the AtaA scaffold and HRP-mediated cross-linking. Increasing the amount of BAP-Y from an AtaA:BAP-Y molar ratio of 1:1 to 1:2 increased the surface-associated enzymatic signal, suggesting that the amount of functionally immobilized BAP-Y can be increased by supplying more BAP-Y during the cross-linking reaction.

To compare the retention of enzymatically active BAP-Y on surfaces coated with the two AtaA constructs, the enzymatic activities of the total reaction mixture, unbound fraction, and surface-bound fraction were measured at an AtaA:BAP-Y molar ratio of 1:1 (Figure 5D). The total BAP activities were comparable between the two systems, indicating that similar amounts of active BAP-Y were initially supplied. However, the YL-NhNs-AtaA system showed higher activity in the unbound fraction (Figure 5D) but lower activity on the solid surface compared with the NhNs-AtaA system (Figure 5E). A direct comparison of the surface-bound fractions confirmed that the enzymatic activity retained on the NhNs-AtaA-coated surface was significantly higher than that retained on the YL-NhNs-AtaA-coated surface (Figure 5E). Comparison of the total and surface-bound activities showed that approximately 1.0 and 0.54% of the initially supplied BAP activity was retained on the NhNs-AtaA- and YL-NhNs-AtaA-coated surfaces, respectively. Although these values indicate that only a small fraction of the supplied enzyme remained functionally active on the surface, the increased surface-associated activity observed at the higher BAP-Y loading (AtaA:BAP-Y molar ratio of 1:2) suggests that the immobilization conditions could be further optimized.

The lower surface-bound activity obtained with YL-NhNs-AtaA is likely attributable to its reduced adhesion to the solid surface, as shown in Figure 1D, rather than to its lower intrinsic ability to undergo tyrosine coupling. YL-NhNs-AtaA showed enhanced intermolecular polymerization in solution, but was likely retained less efficiently on the solid surface before the cross-linking step. Consequently, lower surface-associated BAP activity was obtained despite the higher cross-linking reactivity of the Y-Loop-containing construct. These findings reveal a functional trade-off introduced by the Y-Loop: it promotes protein polymerization but partially compromises the native adhesive function of AtaA. Nevertheless, both constructs enabled the measurable retention of BAP activity on the surface, demonstrating that AtaA can serve as an adhesive scaffold for the covalent immobilization of functional enzymes through HRP-catalyzed tyrosine coupling.

### Potential applications of AtaA-mediated tyrosine coupling

Beyond the present surface-immobilization experiments, the combination of AtaA adhesion and tyrosine coupling may offer several advantages over the previously reported Y-tag- or Y-Loop-based systems. For example, BAP or HRP copolymers containing protein G/A units have been immobilized on microplates using anti-ovalbumin antibodies bound to pre-coated ovalbumin. Thus, these systems require sequential surface coating with an antigen, antibody binding, and the subsequent capture of the enzyme polymer. In contrast, AtaA intrinsically possesses nonspecific adhesiveness toward solid materials, allowing the protein scaffold to be immobilized directly on the surface without prior chemical modification or introduction of a specific ligand–receptor pair. Therefore, the present system simplifies the construction of an enzymatically active surface to direct AtaA adsorption, followed by HRP-mediated coupling of BAP-Y. This self-immobilization property represents a distinct advantage of using AtaA as a scaffold for functional protein coatings.

Another important finding was that NhNs-AtaA lacking the engineered Y-Loop also underwent HRP-mediated tyrosine coupling, indicating that accessible endogenous tyrosine residues in AtaA could function as conjugation sites. Native AtaA is displayed as a long adhesive nanofiber on the bacterial cell surface; therefore, this intrinsic reactivity may allow Y-tagged or Y-Loop-containing enzymes and other functional proteins to covalently attach directly to AtaA-displaying cells. In contrast to conventional protein–polymer immobilization systems that require prior chemical modification or affinity-based functionalization of the target surface, AtaA provides both intrinsic surface adhesion and sites for enzymatic conjugation. We previously developed technologies using AtaA-displaying bacteria ^7,9,11,13,31^ and liposomes ^10^, and the integration of these platforms with tyrosine coupling could enable the construction of multifunctional cell and vesicle surfaces under mild conditions. Although these applications remain to be experimentally validated, the integration of tyrosine coupling with AtaA-based cellular and vesicular platforms may provide a route toward multifunctional biological interfaces under mild conditions.

## Conclusion

This study demonstrated that the adhesive bacterial nanofiber AtaA can serve as a self-immobilizing scaffold for constructing enzymatically active surfaces through HRP-catalyzed tyrosine coupling. Both NhNs-AtaA and YL-NhNs-AtaA formed high-molecular-weight assemblies, indicating that accessible endogenous tyrosine residues in AtaA support intermolecular coupling, whereas the introduction of the Y-Loop further promoted polymerization. The removal of arginine enhanced polymer formation, highlighting the importance of productive intermolecular associations in this reaction. Surface-adsorbed AtaA enabled the functional retention of BAP-Y, yielding catalytically active protein layers without prior chemical or affinity-based modifications of the solid surface. However, the enhanced cross-linking reactivity of YL-NhNs-AtaA was accompanied by reduced adhesion and lower surface-bound phosphatase activity, revealing a trade-off between polymerization and adhesive functions. These findings provide a basis for the direct functionalization of material surfaces and may facilitate the future decoration of AtaA-displaying cells or vesicles for constructing multifunctional interfaces and living materials.

## Experimental Section

### Plasmid construction

Plasmids, primers, and synthetic genes are listed in Supplementary Tables S1–3. NEBuilder HiFi DNA assembly (New England Biolabs, Ipswich, MA, USA) was used to assemble DNA fragments. For the ligation of two DNA fragments, the DNA Ligation Kit Mighty Mix (Takara Bio Inc, Shiga, Japan) was used. To introduce the Y-Loop sequence into the N-terminal domain of an AtaA recombinant protein, the pNhNs plasmid, which carries the gene encording NhNs-AtaA-SNAP-Strep ^10^, was digested using XbaI-EcoRV and ligated with the XbaI-EcoRV-digested fragment of the synthetic gene N-YL-AtaA, generating pNYL-NhNs-AtaA. To introduce the Y-Loop sequence into the C-terminal domain of the AtaA recombinant protein, the pNhNs plasmid was digested with KpnI-NheI and assembled with two PCR fragments, which were amplified using primer sets C-YL3-1_fwd/C-YL3-1_ rev and C-YL3-2_fwd/C-YL3_2_rev from the synthetic gene C-YL-AtaA3 as the DNA template to generate pCYL-NhNs-AtaA3. pNYL-NhNs-AtaA and pCYL-NhNs-AtaA3 were digested with XbaI-EcoRV to construct a recombinant AtaA protein containing both N- and C-terminal Y-Loop sequences. The 1,373 bp-XbaI-EcoRV fragment of pNYL-NhNs and the 10,448 bp-XbaI-EcoRV fragment of pCYL-NhNs were ligated to generate p2YL-NhNs-AtaA3. The synthetic gene fragment BglII-His-Strep-NheI was assembled with BglII-NheI-digested pNCYL-NhNs to generate the pAP05 plasmid. pNhNs and pAP05 were digested with XbaI-NheI. The 8.1 kb-pNhNs fragment and the 3.1 kb-pAP05 fragment were ligated to generate the pAP08 plasmid.

### Bacterial culture

*E. coli* strains were grown in Luria–Bertani or Terrific Broth (TB) medium at 37 °C, with shaking at 200 rpm. Ampicillin (100 µg/mL) was added for the *E. coli* transformants. To produce NhNs-AtaA and YL-NhNs-AtaA recombinant proteins, *E. coli* transformant cells were grown to an optical density at 600 nm (OD_600_) of 0.5; 200 ng/mL anhydrotetracycline (10009542, Cayman Chemical, Ann Arbor, MI) was then added. Subsequently, cells were incubated at 18 °C for 6 h. For the production of BAP-Y recombinant protein, *E. coli* transformant cells harboring pET22b+CY1-BAP(K328R/D330N), which encodes BAP-Y, were grown in TB medium to an OD_600_ of 0.5, and thereafter, 1 mM Isopropyl-β-D-thiogalactopyranoside (19742-94, nacalai tesque, Kyoto, Japan) was added. Cells were incubated at 18 °C for 18 h. The amino acid sequences of the proteins used in this study are listed in the Supporting Information.

### Purification of AtaA recombinant proteins

Both NhNs-AtaA and YL-NhNs-AtaA recombinant proteins were purified using the following procedure: cells were harvested by centrifugation at 8,000 ×*g* for 10 min at 4 °C, resuspended in a buffer (10 mM Tris-HCl, 200 mM arginine-HCl, 5 mM EDTA, 10 µg/mL lysozyme, pH 8.0), lysed by sonication, and centrifuged at 14,000 ×*g* for 30 min at 4 °C. The cell lysate was heated at 60 °C for 30 min, and then centrifuged at 14,000 ×*g* for 30 min at 4 °C. The resultant supernatant was fractionated by precipitation in the 30% solution of the saturated ammonium sulfate at 4 °C. After incubation with stirring for 1 h, the precipitate was collected by centrifugation at 14,000 ×*g* for 15 min at 4 °C and dissolved in 6 M guanidine hydrochloride solution. The concentration of the protein solution was adjusted to less than 1 mg/mL with 6 M guanidine hydrochloride solution, placed in dialysis tubing (2-316-05, Az One Corporation, Tokyo, Japan), and immersed in 250 mL of buffer (10 mM Tris-HCl, 200 mM arginine-HCl, pH 8.0). After incubation with stirring for 24 h at 4 °C, the buffer was exchanged with the same fresh buffer and further incubated for 24 h. Precipitates were removed by filtration using a 0.22 µm membrane, and the filtrate was subjected to size-exclusion chromatography using a HiLoad 16/600 Superdex 200 pg column (Cytiva) with a running buffer (10 mM Tris-HCl, 200 mM arginine-HCl, pH 8.0). Sample purity was confirmed by SDS-PAGE followed by Coomassie Brilliant Blue staining.

### Purification of BAP-Y recombinant protein

BAP-Y recombinant protein was purified following the procedure of a previous study with slight modifications ^32^. Briefly, cells were harvested via centrifugation at 8,000 ×*g* for 10 min at 4 °C, resuspended in a buffer (25 mM Tris-HCl, 137 mM NaCl, 2.68 mM KCl, pH 7.8), lysed by sonication, and centrifuged at 14,000 ×*g* for 30 min at 4 °C. The supernatant was subjected to affinity chromatography using an open column packed with Ni-NTA resin (30210; Qiagen). The desalted elution was subjected to anion-exchange chromatography using a HiTrap Q HP column (Cytiva) with an NaCl gradient of 0–5 M. The fraction containing BAP-Y was purified by size-exclusion chromatography using a HiLoad 16/600 Superdex 200 column with running buffer (50 mM Tris-HCl, 200 mM NaCl, pH 8.0).

### Measurement of CD

The CD spectrum of purified AtaA recombinant proteins (0.1 mg/mL) in 10 mM Tris-HCl (pH 8.0) was recorded using a J-820 spectropolarimeter (JASCO, Tokyo, Japan). The parameters (wavelength range, scan rate, scan interval, response, and bandwidth) were set at 190–250 nm, 100 nm/min, 0.1 nm, 1 s, and 1.0 nm, respectively. The spectrum of 10 mM Tris-HCl buffer (pH 8.0) was recorded as a blank.

### Measurement of adherence

The adhesiveness of purified AtaA recombinant protein was measured using a 27 MHz QCM system (AFFINIX Q8; ULVAC, Kanagawa, Japan). A gold-coated electrode (QCM01S-01; ULVAC) was cleaned with piranha solution (H_2_SO_4_:30% H_2_O_2_ = 3:1) and equilibrated with 99 μL of PBS (140 mM NaCl, 2.7 mM KCl, 10 mM Na_2_HPO_4_, 1.8 mM KH_2_PO_4_, pH 7.4) at 25 °C. After equilibration, 1 μL of 0.1 mg/mL protein sample was added and the dFs were measured at 25 °C for 20 min.

### HRP-catalyzed polymerization of proteins

NhNs-AtaA or YL-NhNs-AtaA was polymerized in a 1.5-mL microcentrifuge tube containing 0.2 mg/mL protein, 3 µM HRP (P8415, Sigma-Aldrich, St. Louis, MO, USA), and 10 µM H₂O₂ in 10 mM Tris-HCl buffer (pH 8.0). The reaction mixture was incubated at 37 °C for 1 h. For copolymerization with BAP-Y, BAP-Y was added at one- or two-fold molar ratios, relative to NhNs-AtaA or YL-NhNs-AtaA. After incubation, the reaction products were analyzed by SDS-PAGE on a 3–14% gradient gel (UH-T/R314; ATTO, Tokyo, Japan). The intensity of the monomeric AtaA band was quantified using ImageJ (Version 1.54 g, National Institutes of Health, Bethesda, MD).

Dityrosine fluorescence was measured using a microplate reader (Synergy HTX; Agilent Technologies) at excitation and emission wavelengths of 320 and 400 nm, respectively. A calibration curve for dityrosine fluorescence was constructed using the same microplate reader (Figure S2). Fluorescence emission spectra were recorded using a spectrofluorometer (FP-6600; JASCO, Tokyo, Japan) at an excitation wavelength of 320 nm.

### Immobilization of copolymer of AtaA and BAP-Y recombinant protein

Seventy-five microliters of protein solution containing 0.35 µM AtaA recombinant protein and equivalent or two-fold molar amount of BAP-Y in the 10 mM Tris-HCl (pH 8.0) was placed in a 96-well plate (196-96FS; WATSON BIO LAB). AtaA recombinant proteins were adhered to the surfaces of wells in a microplate by incubation in a microplate reader at 25 °C for 30 min. Subsequently, the copolymerization reaction was driven by adding 3 µM HRP and 10 µM H_2_O_2_. After the incubation for 1 h at 37 °C, the protein solutions were removed from the wells. To remove unbound proteins, the wells were washed three times with 75 µL of 10 mM Tris-HCl buffer. *p-*NPP was used as the substrate to measure the enzymatic activity of BAP-Y immobilized on the plate.

Seventy-five microliters of 1 mM *p*-NPP solution in 10 mM Tris-HCl was added to the wells, and the absorbance at 410 nm was monitored for 1 h at 37 °C using a microplate reader (Synergy HTX). A calibration curve of *p*-NP absorbance at 410 nm was prepared using the same microplate reader (Figure S3).

### Electron Microscopy

A copper grid with an ultra-high-resolution carbon substrate (UHR-C10; Okenshoji, Tokyo, Japan) was hydrophilized by plasma treatment for 30 min using an HDT-400 (JEOL Ltd., Tokyo, Japan) immediately before use. Ten microliters of the sample solution (3–30 μg/mL) and 2 μl of 50 mM KCl solution were placed onto a copper grid with an ultra-high resolution carbon substrate (UHR-C10; Okenshoji, Tokyo, Japan). The grids were then incubated for 10 min at room temperature to allow adsorption of the proteins and washed by dipping 20 times in 10 mM KCl solution and three times in pure water. Negative staining of the proteins on the grid was performed using the organo-tungstate compound Nano-W (NanoProbes, NY, USA). Specimens were observed using a TEM system (JEM-2100 Plus; JEOL, Tokyo, Japan) operated at 100 kV.

## Author Contributions

**Yuki Yokoe:** Investigation, Formal analysis, Writing – original draft; **Shogo Yoshimoto:** Investigation, Methodology, Writing – review & editing**; Rio Sugimoto:** Investigation, Validation**; Kosuke Minamihata:** Resources, Methodology, Writing – review & editing; **Ayasa Nagatani:** Resources, Methodology, Writing – review & editing; **Atsuki Nara:** Investigation, Writing – review & editing; **Noriho Kamiya:** Resources, Funding acquisition, Writing – review & editing; **Katsutoshi Hori:** Resources, Funding acquisition, Validation, Writing – review & editing; **Masahito Ishikawa:** Resources, Conceptualization, Methodology, Supervision, Validation, Writing – review & editing, Funding acquisition.

## ACKNOWLEDGEMENT

We thank Umechiyo Matsumura, Kazuyo Funatsu, Seiichi Kondo, and Eriko Kawamoto for their technical assistance in this study.

## ABBREVIATIONS

BAP: bacterial alkaline phosphatase
BAP-Y: Y-tagged bacterial alkaline phosphatase
CD: circular dichroism
dF: frequency change
HRP: horseradish peroxidase
p-NP: p-nitrophenol
p-NPP: p-nitrophenyl phosphate
QCM: quartz crystal microbalance
TB: Terrific Broth
TEM: Transmission electron microscopy

