## Supporting Figures and Tables for "AtaA as a Self-Immobilizing Scaffold for Enzymatically Active Surfaces through Oxidative Tyrosine Coupling"

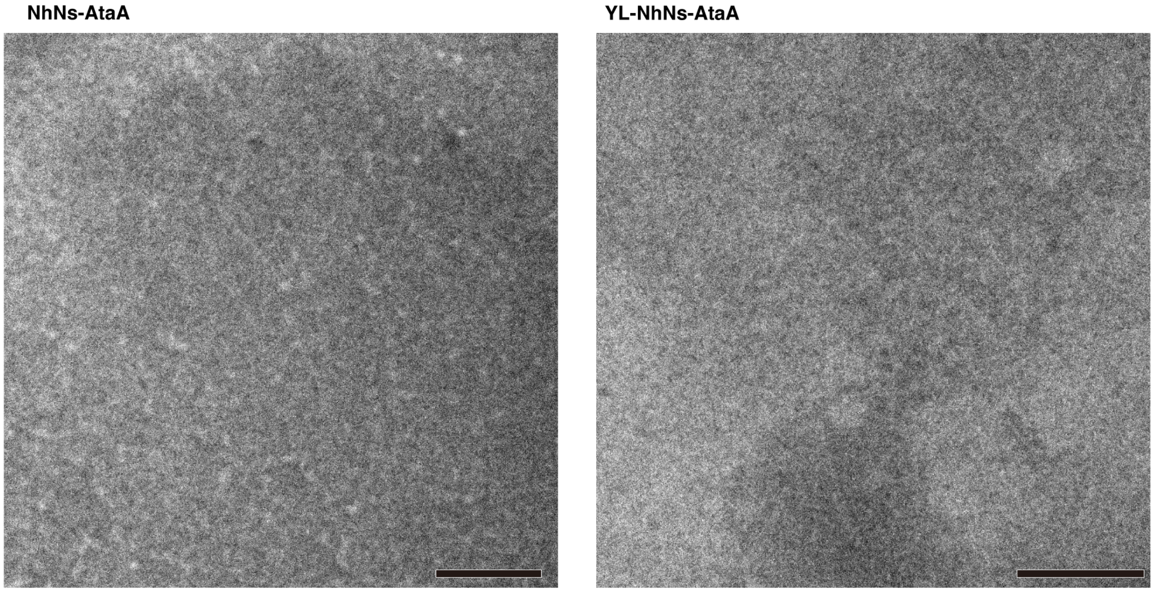


**Figure S1. Transmission electron microscopy images of recombinant AtaA proteins before HRP/$H_2O_2$ treatment.** Representative TEM images of NhNs-AtaA and YL-NhNs-AtaA under arginine-free conditions before HRP/$H_2O_2$ treatment are shown. Scale bars, 100 nm.

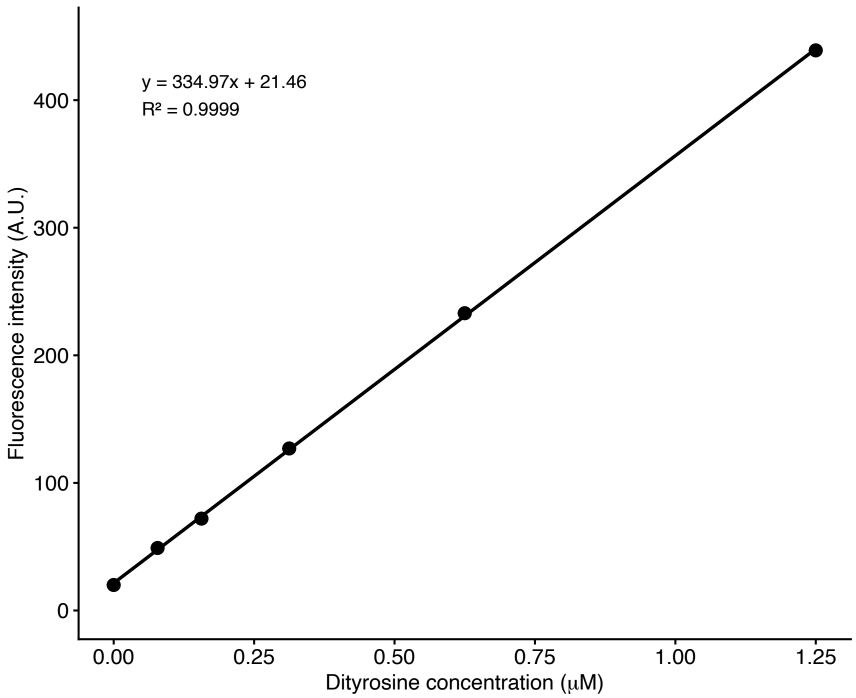


**Figure S2 Calibration curve for dityrosine fluorescence.** Dityrosine fluorescence was measured using a microplate reader at the indicated concentrations. The solid line represents the linear regression fit ($y = 334.97x + 21.46$, $R^2 = 0.9999$).

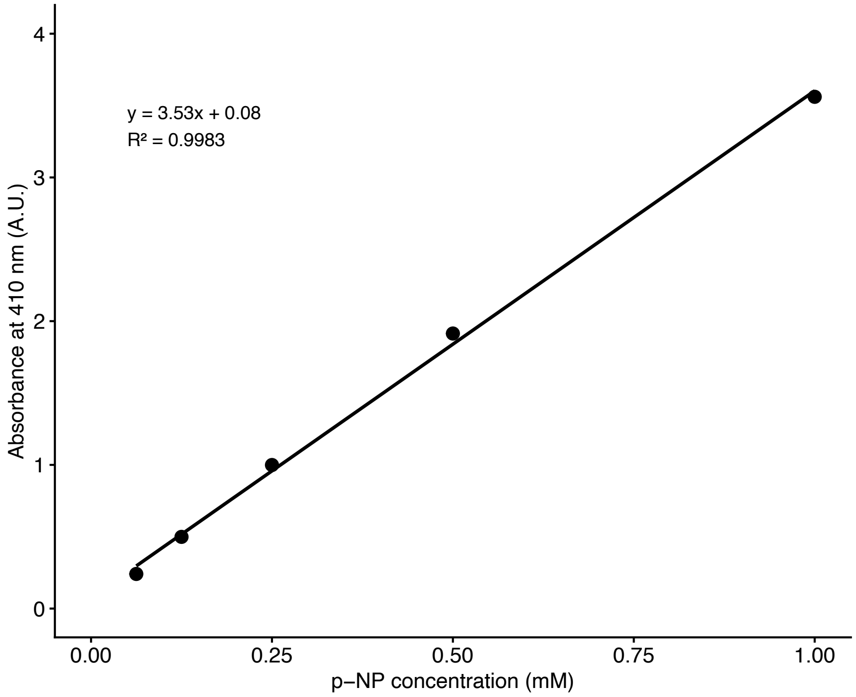


**Figure S3 Calibration curve for *p*-nitrophenol.** The absorbance of *p*-Nitrophenol at 410 nm was measured using a microplate reader at the indicated concentrations. The solid line represents the linear regression fit (y = 3.53x + 0.08, $R^2$ = 0.9983).

**Amino acid sequence of NhNs-AtaA**

MATVQVGGGTNSGTTATASTNCADLYNYQNPENSGSGAAGNYNAGNPSVCSIAIGENAQGG
TSGTGGSPGIAIGGNSKATGGLSVAIGGYAQATNVGSIALGTAALSSGFNSLAISRQAAAT
NNYSIAIGTTSVSKGVGSIAMGHSTNASGDQSIAIGSSDAVNSATATTTYDGTTNTQASGS
KSIAIGASAKASTNNSIALGAGSVTSAQSGNSYLTGVGASATNGVVSVGTSTATRRIQNVA
DGSAASDAVTVAQLDKAYDDTNGRLAAALGTGSGAAYNAANNTYTAPTNIGGTGKNTIDDA
IKATQRSVVAGSNIVVTPTTASDGSISYSVATSATPTFTSITVNNAPTAGTDATNKTYVDS
KAAASRTEVAAGSNVSGVVKTTGANGQDVYTVNANGTTASAGSSAVTVTPGTKDANNVTDY
KVDLSATTKTDIQKGVDAKNAVDTAGLKFKGDTATTSNTKKLGDTVSITGDTNISTVATTD
GVQVKLNPNLDLGATGSVKTGNTTINNAGVTADQVTVGGVVINNTSGINAGGKAITNVAAP
TNNTDAANKKYVDDAGTALTNLGFGLKAQDGTTVNKKLGEAVDIVGSNSNISTKVNAGKVE
VALSNTLDLGTTGSVTTGSTVINNAGVTATQVTANKVTINNAPTAGTDATNKTYVDSKAAA
SRTEVAAGSNVSGVVKTTGANGQDIYAVNANGTTASAGSSAVTVTPGTKDANNVTDYKVDL
SATTKTDIQKGVDAKNAVDTAGLKFKGDTATTSNTKKLGDTVSITGDTNISTVATTDGVQV
KLNPNLDLGATGSVKTGNTTINNAGVTADQVTVGGVVINNTSGINAGGKAITNVAAPTNNT
DAANKKYVDDAGTALTNLGFGLKAQDGTTVNKKLGEAVDIVGSNSNISTKVNAGKVEVALS
NTLDLGTTGSVTTGSTVINNAGVTATQVTANKVTVNNAPTAGTDATNKTYVDSKAAASRTE
VAAGSNVSGVVKTTGANGQDVYTVNANGTTASAGSSAVTVTPGTKDANNVTDYKVDLSATT
KTDIQKGVDAKNAVDTAGLKFKGDTATTSNTKKLGDTVSITGDTNISTVATTDGVQVKLNP
NLDLGATGSVKTGNTTINNAGVTADQVTVGGVVINNTSGINAGGKAITNVAAPTNNTDAAN
KKYVDDAGTALTNLGFGLKAQDGTTVNKKLGEAVEVVGADSNITTKVAGGQVAIELNKNLN
NLTGITVNDGTNGTNGSTVIGKDGISVKDGSGNTIAGVDNTALTVKDGSGNTETSINQAIN
TLNAAQGETDKFAVKYDKNADGSVNYNNITLAGTTASSTQDATTGKITTTGGTSLNNVASA
GDYKDVANASKGVNAGDLNNAVVDATNAATSKGFALQAADGAKVQKNLGEAVEVVGADSNI
TTKVAGGQVAIELNKNLNNLTGITVNDGTNGTNGSTVIGKDGISVKDGSGNTIAGVDNTAL
TVKDGSGNTETSINQAINTLNAAQGETDKFAVKYDKNTDGSTNYNSITAGNGNGTAATIGT
DTAGNSVVTSGGTKISNVANGVNASDAVNKGQLDSLSTGLTNTGFGLKAADGNTVNKKLGE
AVDVVGADSNITTKVAGGQVAIELNKNLNNLTGITVNDGTNGTNGSTVIGKDGISIKDGSG
NTIAGVDNTALTVKDGSGNTETSINQAINTLNAAQGETDKFAVKYDKNADGSANYNNITLA
GTTASSTQDATTGKITTTGGTSLNNVASAGDYKDVANASKGVNAGDLNNAVVDATNAATSK
GFALQAADGAKVQKNLGEAVEVVGADSNITTKVVGGQVAIELNKNLNNLTGITVNDGTNGT
NGSTVIGKDGISVKDGSGNTIAGVDNTALTVKDGSGNTETSINQAINTLNAAQGETDKFAV
KYDKNADGSVNYNNITLAGTTASSTQDATTGKITTTGGTSLNNVASAGDYKDVANASKGVN
AGDLNNAVVDATNAATSKGFALQAADGAKVQKNLGEAVEVVGADSNITTKVAGGQVAIELN
KNLNNLTGITVNDGTNGTNGSTVIGKDGISVKDGSGNTIAGVDNTALTVKDGSGNTETSIN
QAINTLNAAQGETDKFAVKYDKNADGSVNYNNITLAGTTASSTQDATTGKITTTGGTSLNN
VASAGDYKDVANASKGVNAGDLNNAVVDATNAATSKGFALQAADGAKVQKNLGEAVEVVGA
DSNITTKVAGGQVAIELNKNLNNLTGITVNDGTNGTNGSTVIGKDGISVKDGSGNTIAGVD
NTALTVKDGSGNTETSINQAINTLNAAQGETDKFAVKYDKNADGSANYNNVTLAGTNGTII
SNVKAGAVTSTSTDAINGSQLYGVANSVKNAIGGSTTIDATTGAITTTNIGGTGSNTIDGA
ISSIKDSATKAKTTVSAGDNVVVTSGTNADGSTNYEVATAKDVNFDKVTVGSVVVDKSSNT
IKGLSNTTWNGTAVSGQAATEDQLKTVSDAQGETDKFAVKYDKNADGSANYNSITAGNGNG
TAATIGTDTAGNSVVTSGGTKISNVANGVNASDAVNKGQLDSLSTGLTNTGFGLKAADGNT
VNKKLGEAVDVVGADSNITTKVAGGQVAIELNKNLNNLTGITVNDGTNGTNGSTVIGKDGI

SIKDGSGNTIAGVDNTALTVKDSSGNTETSINQAINTLNAAQGETDKFAVKYDKNADGSVN
YNNVTLAGTNGTIIRNVKAGAVTSTSTDAINGSQLYDIANSVKNAIGGSTTRDVTTGAITT
TNIGGTGSNTIDGAISSIKDSATKAKTTISAGDNVVVTSGTNADGSTNYEVATAKDVNFDK
VTVGNVVVDKANDTIQGLSNKDLNSTDFATKGRAATEEQLKQIEDKIEEILSKIYHIENEI
ARIKKLIKLHHHHHHGGGSAWSHPQFEK

**Amino acid sequence of YL-NhNs-AtaA**

MATVQVGGGTNSGTTATASTNCADLYNYQNPENSGSGAAGNYNAGNPSVCSIAIGENAQGG
TSGTGGSPGIAIGGNSKATGGLSVAIGGYAQATNVGSIALGTAALSSGFNSLAISRQAAAT
NNYSIAIGTTSVSKGVGSIAMGHSTNASGDQSIAIGSSDAVNSATATTTYDGTTNTQASGS
KSIAIGASAKASTNNSIALGAGSVTSAQSGNSYLTGVGASATNGVVSVGTSTATRRIQNVA
DGSAASDAVTVAQLDKAYDDTNGRLAAALIRINRGPGYAFVTGTGSGAAYNAANNTYTAPT
NIGGTGKNTIDDAIKATQRSVVAGSNIVVTPTTASDGSISYSVATSATPTFTSITVNNAPT
AGTDATNKTYVDSKAAASRTEVAAGSNVSGVVKTTGANGQDVYTVNANGTTASAGSSAVTV
TPGTKDANNVTDYKVDLSATTKTDIQKGVDAKNAVDTAGLKFKGDTATTSNTKKLGDTVSI
TGDTNISTVATTDGVQVKLNPNLDLGATGSVKTGNTTINNAGVTADQVTVGGVVINNTSGI
NAGGKAITNVAAPTNNTDAANKKYVDDAGTALTNLGFGLKAQDGTTVNKKLGEAVDIVGSN
SNISTKVNAGKVEVALSNTLDLGTTGSVTTGSTVINNAGVTATQVTANKVTINNAPTAGTD
ATNKTYVDSKAAASRTEVAAGSNVSGVVKTTGANGQDIYAVNANGTTASAGSSAVTVTPGT
KDANNVTDYKVDLSATTKTDIQKGVDAKNAVDTAGLKFKGDTATTSNTKKLGDTVSITGDT
NISTVATTDGVQVKLNPNLDLGATGSVKTGNTTINNAGVTADQVTVGGVVINNTSGINAGG
KAITNVAAPTNNTDAANKKYVDDAGTALTNLGFGLKAQDGTTVNKKLGEAVDIVGSNSNIS
TKVNAGKVEVALSNTLDLGTTGSVTTGSTVINNAGVTATQVTANKVTVNNAPTAGTDATNK
TYVDSKAAASRTEVAAGSNVSGVVKTTGANGQDVYTVNANGTTASAGSSAVTVTPGTKDAN
NVTDYKVDLSATTKTDIQKGVDAKNAVDTAGLKFKGDTATTSNTKKLGDTVSITGDTNIST
VATTDGVQVKLNPNLDLGATGSVKTGNTTINNAGVTADQVTVGGVVINNTSGINAGGKAIT
NVAAPTNNTDAANKKYVDDAGTALTNLGFGLKAQDGTTVNKKLGEAVEVVGADSNITTKVA
GGQVAIELNKNLNNLTGITVNDGTNGTNGSTVIGKDGISVKDGSGNTIAGVDNTALTVKDG
SGNTETSINQAINTLNAAQGETDKFAVKYDKNADGSVNYNNITLAGTTASSTQDATTGKIT
TTGGTSLNNVASAGDYKDVANASKGVNAGDLNNAVVDATNAATSKGFALQAADGAKVQKNL
GEAVEVVGADSNITTKVAGGQVAIELNKNLNNLTGITVNDGTNGTNGSTVIGKDGISVKDG
SGNTIAGVDNTALTVKDGSGNTETSINQAINTLNAAQGETDKFAVKYDKNTDGSTNYNSIT
AGNGNGTAATIGTDTAGNSVVTSGGTKISNVANGVNASDAVNKGQLDSLSTGLTNTGFGLK
AADGNTVNKKLGEAVDVVGADSNITTKVAGGQVAIELNKNLNNLTGITVNDGTNGTNGSTV
IGKDGISIKDGSGNTIAGVDNTALTVKDGSGNTETSINQAINTLNAAQGETDKFAVKYDKN
ADGSANYNNITLAGTTASSTQDATTGKITTTGGTSLNNVASAGDYKDVANASKGVNAGDLN
NAVVDATNAATSKGFALQAADGAKVQKNLGEAVEVVGADSNITTKVVGGQVAIELNKNLNN
LTGITVNDGTNGTNGSTVIGKDGISVKDGSGNTIAGVDNTALTVKDGSGNTETSINQAINT
LNAAQGETDKFAVKYDKNADGSVNYNNITLAGTTASSTQDATTGKITTTGGTSLNNVASAG
DYKDVANASKGVNAGDLNNAVVDATNAATSKGFALQAADGAKVQKNLGEAVEVVGADSNIT
TKVAGGQVAIELNKNLNNLTGITVNDGTNGTNGSTVIGKDGISVKDGSGNTIAGVDNTALT
VKDGSGNTETSINQAINTLNAAQGETDKFAVKYDKNADGSVNYNNITLAGTTASSTQDATT
GKITTTGGTSLNNVASAGDYKDVANASKGVNAGDLNNAVVDATNAATSKGFALQAADGAKV
QKNLGEAVEVVGADSNITTKVAGGQVAIELNKNLNNLTGITVNDGTNGTNGSTVIGKDGIS

VKDGSGNTIAGVDNTALTVKDGSGNTETSINQAINTLNAAQGETDKFAVKYDKNADGSANY
NNVTLAGTNGTIISNVKAGAVTSTSTDAINGSQLYGVANSVKNAIGGSTTIDATTGAITTT
NIGGTGSNTIDGAISSIKDSATKAKTTVSAGDNVVVTSGTNADGSTNYEVATAKDVNFDKV
TVGSVVVDKSSNTIKGLSNTTWNGTAVSGQAATEDQLKTVSDAQGETDKFAVKYDKNADGS
ANYNSITAGNGNGTAATIGTDTAGNSVVTSGGTKISNVANGVNASDAVNKGQLDSLSTGLT
NTGFGLKAADGNTVNKKLGEAVDVVGADSNITTKVAGGQVAIELNKNLNNLTGITVNDGTN
GTNGSTVIGKDGISIKDGSGNTIAGVDNTALTVKDSSGNTETSINQAINTLNAAQGETDKF
AVKYDKNADGSVNYNNVTLAGTNGTIIRNVKAGAVTSTSTDAINGSQLYDIANSVKNAIIR
INRGPGYAFVTGGSTTRDVTTGAITTTNIGGTGSNTIDGAISSIKDSATKAKTTISAGDNV
VVTSGTNADGSTNYEVATAKDVNFDKVTVGNVVVDKANDTIQGLSNKDLNSTDFATKGRAA
TEEQLKQIEDKIEEILSKIYHIENEIARIKKLIKLHHHHHHGGGSAWSHPQFEK

**Amino acid sequence of BAP-Y**

MKYLLPTAAAGLLLLAAQPAMA↓MDIGINSDPHHHHHHTPEMPVLENRAAQGDITAPGGAR
RLTGDQTAALRDSLSDKPAKNIILLIGDGMGDSEITAARNYAEGAGGFFKGIDALPLTGQY
THYALNKKTGKPDYVTDSAASATAWSTGVKTYNGALGVDIHEKDHPTILEMAKAAGLATGN
VSTAELQDATPAALVAHVTSRKCYGPSATSEKCPGNALEKGGKGSITEQLLNARADVTLGG
GAKTFAETATAGEWQGKTLREQAQARGYQLVSDAASLNSVTEANQQKPLLGLFADGNMPVR
WLGPKATYHGNIDKPAVTCTPNPQRNDSVPTLAQMTDKAIELLSKNEKGFFLQVEGASIDR
QNHAANPCGQIGETVDLDEAVQRALEFAKKEGNTLVIVTADHAHASQIVAPDTKAPGLTQA
LNTKDGAVMVMSYGNSEEDSQEHTGSQLRIAAYGPHAANVVGLTDQTDLFYTMKAALGLKL
VPRGSGGGGY

Magenta: Y-Loop
Orange: GCN4 adaptor
Green: His-tag
Purple: Strep-Tag II
Red: Signal peptide
Underline: Mutation for improving enzymatic activity
Grey: Thrombin recognition sequence
Blue: Y-tag

**Table S1. Plasmids in this study**

| Plasmid | Description | Reference |
|---|---|---|
| pNhNs | Expression plasmid encoding AtaA-SNAP-Strep | [1] |
| pNYL-NhNs-AtaA | pNhNs derivative encoding AtaA-SNAP-Strep with an N-terminal Y-Loop | This study |
| pCYL-NhNs-AtaA3 | pNhNs derivative encoding AtaA-SNAP-Strep with an C-terminal Y-Loop | This study |
| p2YL-NhNs-AtaA3 | pNhNs derivative encoding AtaA-SNAP-Strep with N- and C-terminal Y-Loop | This study |
| pAP05 | Expression plasmid encoding YL-NhNs-AtaA | This study |
| pAP08 | Expression plasmid encoding NhNs-AtaA | This study |
| pET22b+CY1-BAP(K328R/D330N) | pET-22b encoding recombinant BAP that includes N-terminal His-tag, two mutations (K328R/D330N), and C-terminal Y-tag. | This study |

**Table S2. Synthetic genes used in this study**

| Synthetic gene | Sequence (5'→3') | Vender |
|---|---|---|
| N-YL-AtaA | ATGAATAGTTCGACAAAAATCTAGATAACGAGGGCAAAAAATGG<br>CTACTGTTCAAGTTGGAGGGGGAACTAATTCTGGAACAACTGCT<br>ACAGCTTCTACGAATTGTGCAGACTTATATAATTATCAAAATCC<br>TGAGAACTCAGGCTCTGGAGCGGCTGGGAATTATAATGCAGGAA<br>ATCCAAGTGTGTGTTCGATCGCTATAGGTGAAAACGCACAAGGT<br>GGTACTTCTGGAACTGGAGGGTCGCCAGGGATAGCGATAGGTGG<br>AAATTCTAAAGCTACGGGTGGTTTATCTGTTGCTATAGGCGGAT<br>ATGCTCAAGCGACAAATGTTGGAAGTATTGCTTTAGGCACAGCA<br>GCTTTATCAAGTGGTTTTAACAGTTTAGCAATATCCAGACAAGC<br>TGCTGCAACGAATAACTATTCAATAGCTATAGGTACAACTTCAG<br>TTTCGAAAGGAGTTGGATCGATTGCTATGGGGCATTCAACGAAT<br>GCTTCTGGAGATCAATCGATAGCAATTGGTAGCTCGGATGCTGT<br>TAATTCAGCAACAGCAACAACAACATACGATGGTACAACAAATA<br>CTCAAGCATCAGGTAGTAAATCGATTGCTATAGGTGCAAGCGCA<br>AAGGCATCAACCAATAACAGCATTGCACTAGGTGCAGGATCGGT<br>AACTTCTGCACAATCTGGTAATTCTTATCTTACTGGTGTAGGTG<br>CATCAGCTACAAATGGTGTTGTATCTGTTGGAACTTCAACTGCA<br>ACACGTCGTATCCAAAATGTAGCAGATGGTTCAGCCGCTTCAGA<br>TGCTGTGACAGTTGCTCAGTTGGATAAAGCTTATGATGATACAA<br>ATGGTCGTTTAGCTGCTGCTTTAATCCGTATCAACCGTGGTCCG<br>GGTTACGCTTTCGTTACTGGTACAGGTAGTGGTGCTGCCTATAA<br>TGCAGCAAACAATACATATACCGCTCCAACGAATATTGGGGGAA<br>CAGGTAAAAATACGATTGATGATGCAATTAAAGCAACTCAACGA<br>AGTGTAGTCGCTGGATCAAATATTGTCGTTACCCCGACGACAGC<br>TTCTGATGGTTCAATATCGTATTCGGTTGCTACAAGCGCAACAC<br>CGACGTTTACAAGTATAACTGTAAACAATGCACCAACGGCAGGT<br>ACAGATGCGACCAACAAGACTTATGTAGACTCAAAAGCAGCAGC<br>ATCGAGAACAGAAGTAGCAGCTGGAAGCAATGTATCTGGTGTAG<br>TAAAAACGACAGGCGCAAACGGTCAAGACGTTTATACAGTAAAT<br>GCCAATGGTACGACTGCATCAGCAGGTTCTTCAGCAGTTACCGT<br>AACACCAGGCACGAAAGATGCAAATAATGTCACTGACTATAAAG<br>TAGACTTATCAGCGACTACAAAAACCGATATCCAAAAAGGTGTA<br>GATGCAAAGATATCCAAAAAGGTGTAGATGCAAA | GenScript |
| C-YL-AtaA3 | ATAATGTTGTCGTTACATCGGGTACCAATGCAGATGGCTCAACA<br>AACTATGAAGTTGCGACAGCGAAAGACGTTAACTTTGACAAAGT<br>GACTGTAGGTAGTGTTGTTGTAGATAAATCAAGCAATACAATCA<br>AAGGATTAAGTAATACCACTTGGAACGGAACAGCAGTATCAGGT<br>CAAGCGGCGACAGAAGACCAGTTAAAAACGGTCAGCGATGCGCA<br>AGGTGAAACTGATAAGTTTGCAGTGAAGTACGACAAAAATGCTG<br>ATGGCAGTGCGAACTACAACAGTATTACTGCAGGCAATGGTAAC<br>GGTACTGCAGCAACGATCGGAACTGACACAGCAGGTAATAGTGT<br>TGTGACCAGTGGCGGAACTAAAATTAGTAATGTTGCGAATGGTG<br>TCAATGCAAGTGATGCAGTAAACAAAGGTCAATTGGATAGCTTA<br>AGTACAGGTCTTACCAATACAGGCTTTGGTTTAAAAGCAGCAGA<br>TGGCAACACCGTTAACAAAAAATTAGGCGAAGCAGTAGACGTTG<br>TCGGTGCTGACAGCAACATCACCACGAAAGTTGCAGGCGGTCAG | Genscript |

| | | |
|---|---|---|
| | GTTGCGATTGAGTTAAATAAAAACCTCAACAACTTAACTGGCAT<br>TACCGTGAACGATGGAACCAATGGCACCAATGGTTCAACTGTGA<br>TTGGTAAAGATGGTATTTCGATTAAAGATGGTTCAGGCAATACC<br>ATTGCAGGTGTAGATAACACAGCGTTGACGGTTAAAGATAGCAG<br>TGGCAACACAGAAACCAGCATTAACCAAGCGATCAACACGTTAA<br>ATGCAGCGCAAGGTGAAACTGATAAGTTTGCAGTGAAGTACGAT<br>AAGAATGCTGATGGCAGTGTGAACTATAACAATGTCACTTTAGC<br>AGGTACAAATGGCACAATAATCAGAAATGTTAAAGCGGGTGCTG<br>TGACCTCAACATCTACTGATGCGATCAATGGTAGCCAATTATAC<br>GATATTGCAAACAGCGTGAAGAATGCAATTATCCGTATCAACCG<br>TGGTCCGGGTTACGCTTTCGTTACTGGTGGTTCAACCACAAGAG<br>ATGTAACGACTGGTGCAATCACAACGACCAATATTGGTGGTACA<br>GGTTCAAACACGATTGATGGTGCAATCAGCAGTATTAAAGATTC<br>AGCGACTAAAGCGAAAACCACGATAAGTGCTGGGGATAATGTTG<br>TCGTTACATCGGGTACCAATGCAGATGGCTCAACAAACTATGAA<br>GTTGCGACAGCGAAAGACGTTAACTTTGACAAAGTAACTGTAGG<br>TAATGTTGTTGTTGATAAGGCAAATGACACGATCCAAGGTTTGA<br>GCAATAAAGATCTAAATTCAACTGATTTTGCGACCAAAGGTAGA<br>GCTGCGACTGAAGAACAGTTAAAACAGATTGAAGATAAAATTGA<br>AGAAATTCTGAGCAAAATTTATCATATTGAAAACGAAATTGCGC<br>GTATTAAAAAACTGATTAAGCTTTGAGCTAGGCTAGCAGCTTGA<br>AAAATGGCGCAC | |
| BglII-His-Strep-NheI | TCCAAGGTTTGAGCAATAAAGATCTAAATTCAACTGATTTTGCG<br>ACCAAAGGTAGAGCTGCGACTGAAGAACAGTTAAAACAGATTGA<br>AGATAAAATTGAAGAAATTCTGAGCAAAATTTATCATATTGAAA<br>ACGAAATTGCGCGTATTAAAAAACTGATTAAGCTTCACCACCAC<br>CACCACCACGGCGGCGGCAGCGCCTGGAGCCACCCGCAGTTCGA<br>AAAATGAGCTAGCAGCTTGAAAAATGGCGCAC | IDT |

**Table S3. Primers used in this study**

| Primer | Sequence (5'→3') |
|---|---|
| C-YL3-1_fwd | ATAATGTTGTCGTTACATCGG |
| C-YL3-1_rev | TGCTGGTTTCTGTGTTGCCACTGCTATC |
| C-YL3-2_fwd | TGGCAACACAGAAACCAGCATTAACCAAG |
| C-YL3-2_rev | GTGCGCCATTTTTCAAGC |
